# Inhibitory control in reptiles

**DOI:** 10.1101/2025.02.12.637795

**Authors:** Maria Santacà, Anna Wilkinson, Gionata Stancher, Valeria Anna Sovrano, Angelo Bisazza

## Abstract

Inhibitory control, the ability to suppress an automatic response in favour of a contextually appropriate alternative, is crucial for adaptive behaviour across animal species. While extensively studied in mammals and birds, research on reptiles remains limited, with comparisons hindered by methodological inconsistencies. Here, we assessed inhibitory control in two reptile species, Hermann’s tortoise (*Testudo hermanni*) and the bearded dragon (*Pogona vitticeps*), using the transparent cylinder test—a method widely employed with mammals, birds, and fish. This test evaluates the ability to inhibit reaching directly for visible food through a transparent barrier. Both species exhibited lower inhibitory control than most amniotes, supporting the prevalent hypothesis linking inhibitory capacity to brain size. However, exceptions observed in various species suggest ecological and non-cognitive factors also shape these abilities. Notably, bearded dragons outperformed tortoises, despite their smaller size. In tortoises, females surpassed males, highlighting sex-based inhibitory differences in non-avian reptiles. Additionally, a positive correlation between lateralization and inhibitory control was observed in both species, providing the first evidence of such a link in reptiles. These findings emphasize the role of lateralization in reptilian cognition and suggest that inhibitory control across vertebrates is influenced by diverse factors, including brain size, ecology, and sex.

## INTRODUCTION

Inhibitory control refers to the capacity to suppress a prepotent or automatic response in favour of a more context-appropriate alternative, playing a crucial role in adaptive behaviours. For instance, ambush predators delay their attack until prey is optimally positioned [1], while prey species restrain feeding in the presence of threats [2]. In hierarchically structured primate societies, subordinates inhibit feeding or sexual responses in the presence of higher-ranking individuals [3].

Inhibitory capacities have been studied in a wide variety of organisms. While some studies have been conducted in natural settings [4,5], most research on inhibitory control in animals has been performed in the laboratory. The transparent cylinder test is a benchmark method for studying inhibition. It can be easily administered to diverse species, allowing direct comparisons even between phylogenetically distant animal species. In this test, an individual is first trained to retrieve a food reward from an opaque cylinder that is open at both ends. Once it reliably performs this task, the opaque cylinder is replaced with a transparent one. The test evaluates the individual’s ability to inhibit the motor impulse to reach directly for the reward through the side of the cylinder, which acts as a transparent barrier, and instead navigate to the open ends to obtain the reward.

Historically, research has focused on mammals and birds, revealing considerable interspecific variation in inhibitory capacity, with certain groups, such as primates and corvids, outperforming other groups [6,7]. A large-scale, multi-lab study on 36 mammal and bird species found that the best predictor of performance was absolute brain volume [6]. Inhibitory capacity also correlated with general intelligence (or g-factor), and complex cognitive abilities such as innovation, tool use, and social learning. They suggested that the reason inhibitory control correlates with brain mass is that larger brains have greater computational capacity. However, brain mass alone is only a loose predictor of a species’ information processing capacity, as neuronal density varies significantly across taxa [8]. Herculano-Houzel [9] re-analysed the data sets from MacLean and collaborators [6], along with a subsequent study on three corvid species [7], identifying the total number of cortical or pallial neurons as a stronger predictor of cylinder test performance. Additionally, ecological factors, such as diet, may further influence inhibitory capacities [6]. Species with broader diets tend to exhibit more advanced cognitive functions, as they face complex decision-making in varied foraging contexts. This may partially explain the superior performance observed in omnivorous species like corvids and terrestrial primates.

More recently, inhibitory control has been investigated in teleost fish [10,11]. Remarkably, despite their smaller brains, teleosts exhibit inhibitory abilities comparable to endotherms. This suggests the relationship with brain size might apply only to endotherms. Alternatively, as indicated by other research [13–17], the cognitive abilities of teleosts might be distinct from the other vertebrates due to the long, independent radiation and evolution of this group following their separation from the lineage that gave origin to terrestrial tetrapods. They might for instance differ in connectivity, nonneuronal/neuronal cell ratio or neuronal density [12].

To resolve this issue, data from a wider range of vertebrates are needed. No studies exist on cartilaginous fish or amphibians, and only one recent study has examined reptiles. This study assessed five skink species, finding differences in motor inhibition, with blue-tongued skinks making fewer errors than all other species [18]. While this study provides valuable insights and highlights interspecies differences, it is unsuitable for interspecific comparison due to methodological differences, primarily because the study used a semi-transparent mesh cylinder rather than a fully transparent one. Transparency levels are known to have a significant impact on such tests [19–21], making direct comparisons with the extensive work on other species challenging. In addition to inhibitory control, the study explored reversal learning [18], which tests cognitive flexibility by requiring the inhibition of a previously reinforced response and adaptation to a new rule. Unlike inhibitory control, reversal learning assesses the ability to adjust to changing stimulus-response associations rather than suppress impulsive actions. Szabo et al. [18] found no correlation between cylinder task performance and reversal learning in skinks, suggesting that these processes may rely on different neural mechanisms or that the methodology used may have influenced the results. Unlike the cylinder task, which involves resisting a direct, prepotent response, the reversal learning task requires suppressing a previously reinforced choice. The observed lack of association might indicate that these two abilities are not strongly linked in reptiles, potentially due to differing ecological pressures shaping each skill. Alternatively, it could suggest that the two tasks do not capture overlapping cognitive demands.

Cerebral lateralization - the specialization of left and right hemispheres for different cognitive functions - is widespread across all vertebrate classes [22]. In fish and endotherms, lateralized individuals have been shown to be advantaged in a range of cognitive functions [23,24]. Rogers [22,25] suggested that hemispheric specialization offers a selective advantage by enabling more efficient simultaneous task performance, a hypothesis supported by substantial empirical evidence [23,26,27]. Recent studies have further identified a correlation between the direction of lateralization and inhibitory capacity [28,29].

Regarding the two species investigated in this study, previous research on bearded dragons (*Pogona vitticeps*) showed a consistent eye preference when observing stimuli, suggesting hemispheric specialization in visual processing [30,31]. Similarly, in Hermann’s tortoises (*Testudo hermanni*), lateralized behaviour has been observed in righting reflexes, with a preference for turning in one direction to regain an upright position [32], as well as in interactions with mirrors [33]. These studies underscore the relevance of lateralization in diverse behavioural contexts, raising the possibility that in our two species it might also influence inhibitory control performance.

This research aims to investigate inhibitory control in reptiles using the transparent cylinder test, the method previously applied to mammals, birds, and fish. To this end, we selected two phylogenetically distant species: a tortoise (*Testudo hermanni*) and an iguanid lizard (*Pogona vitticeps*). In both species, we measured individual lateralization to assess whether this phenomenon also affects inhibitory performance. For tortoises, which had a larger sample size, we also examined sex differences.

## MATERIALS AND METHODS

### Ethical note

This study adhered strictly to ethical standards for animal research, ensuring that both tortoises (*Testudo hermanni*) and bearded dragons (*Pogona vitticeps*) experienced minimal stress throughout housing, testing, and post-experiment care. Animal husbandry and experimental procedures complied with European Legislation for the Protection of Animals (Directive 2010/63/ EU) and the ARRIVE guidelines. The study with tortoises was carried out at the natural estate of “SperimentArea” (Civic Museum Foundation of Rovereto, Trento, Italy) and was done in accordance with the Italian and European Community laws on protected wild species (Art. 8/bis 150/92 all. A Reg. (CE) 338/97). The experimental protocol was authorized by the internal Ethics Committee of the Civic Museum Foundation of Rovereto (Prot. A N. 0000238 – dd 21/06/23). The study with bearded dragons was carried out at the University of Lincoln. Applicable national guidelines for the care and use of animals were followed. This research was approved by the ethics committee of the School of Life Sciences, University of Lincoln (protocol N. CoSREC364).

Animals were housed in enriched environments suited to their species-specific needs before, during and after the experiment (see below for details). For tortoises, testing was scheduled during the animals’ peak activity season, further reducing stress associated with temperature or seasonal inactivity. The testing protocol, the transparent cylinder task, relies on spontaneous observation of behaviour, with all our subjects participating voluntarily and showing no signs of stress. In each trial, animals were provided with a preferred food reward and received additional feeding post-trial. Measures were taken to minimize the number of animals used, following the principle of “Reduction”, to observe the minimum number of animals to draw statistically valid results. At the conclusion of the experiments, all animals were returned to their original enclosures, ensuring their well-being and continuity of care.

### Subjects

Experiments were conducted on a sample of 24 adult tortoises (*Testudo hermanni*, 12 males and 12 females) and 8 adult bearded dragons (*Pogona vitticeps*, 6 females and 2 males). Tortoises were maintained in groups in an open-air enclosure at the SperimentArea, which is under the supervision of the Rovereto Civic Museum Foundation, located in Italy. During the study, the tortoises had unrestricted access to clean water and shelters. The experiments were run during the period spanning from June to July, corresponding to the months of higher activity of this species. We matched body size between the sexes as much as possible; indeed, we found no significant difference in their size considering the Straight Carapace Length (SCL): one-sample *t*-test, *t*_22_ = - 0.086, *p* = 0.932. Bearded dragons were housed individually or in pairs in vivaria at the School of Life Sciences, University of Lincoln. During the study, the bearded dragons had full unrestricted access to clean water, shelter, UV light and heat lamps. All reptiles were not food deprived during the experiments; they received a favoured food reward during the experiment and were also fed after finishing the daily trials.

### Stimuli and apparatus

The stimuli consisted of small pieces of tomatoes and red cabbages for the tortoises and vegetable extract (kale, cucumber and mint) jellies for bearded dragons, highly preferred food for both species. Vegetables and jellies were freshly prepared each day.

For bearded dragons, the apparatus consisted of a rectangular arena (110□×□61□cm; Figure 1a) entirely covered with black plastic and was placed in an illuminated test room. For tortoises, the apparatus consisted of a wooden arena (90□×□90□cm; Figure 1b) that was centrally lit from above (∼150 cm) through a 400 W halogen lamp. The arena was placed in a dark room. The inner part of the arena was covered with wooden pellets. All tests were video recorded with a video camera placed above the apparatuses. For all reptiles, the cylinder test took place near an apparatus wall, opposite to where they were inserted in the test apparatus. We used two types of cylinders in the two different phases of the procedure: the training cylinder was opaque (green plastic), whereas the test cylinder was transparent (acetate). For bearded dragons, they were both 15 cm in length and 10 cm in diameter (Figure 1a) whereas, for tortoises, they were both 30 cm in length and 20 cm in diameter (Figure 1b). All cylinders for both species were glued onto a plastic base (tortoises: 30 × 30 cm; bearded dragons: 15 × 15 cm).

**Figure 1.**
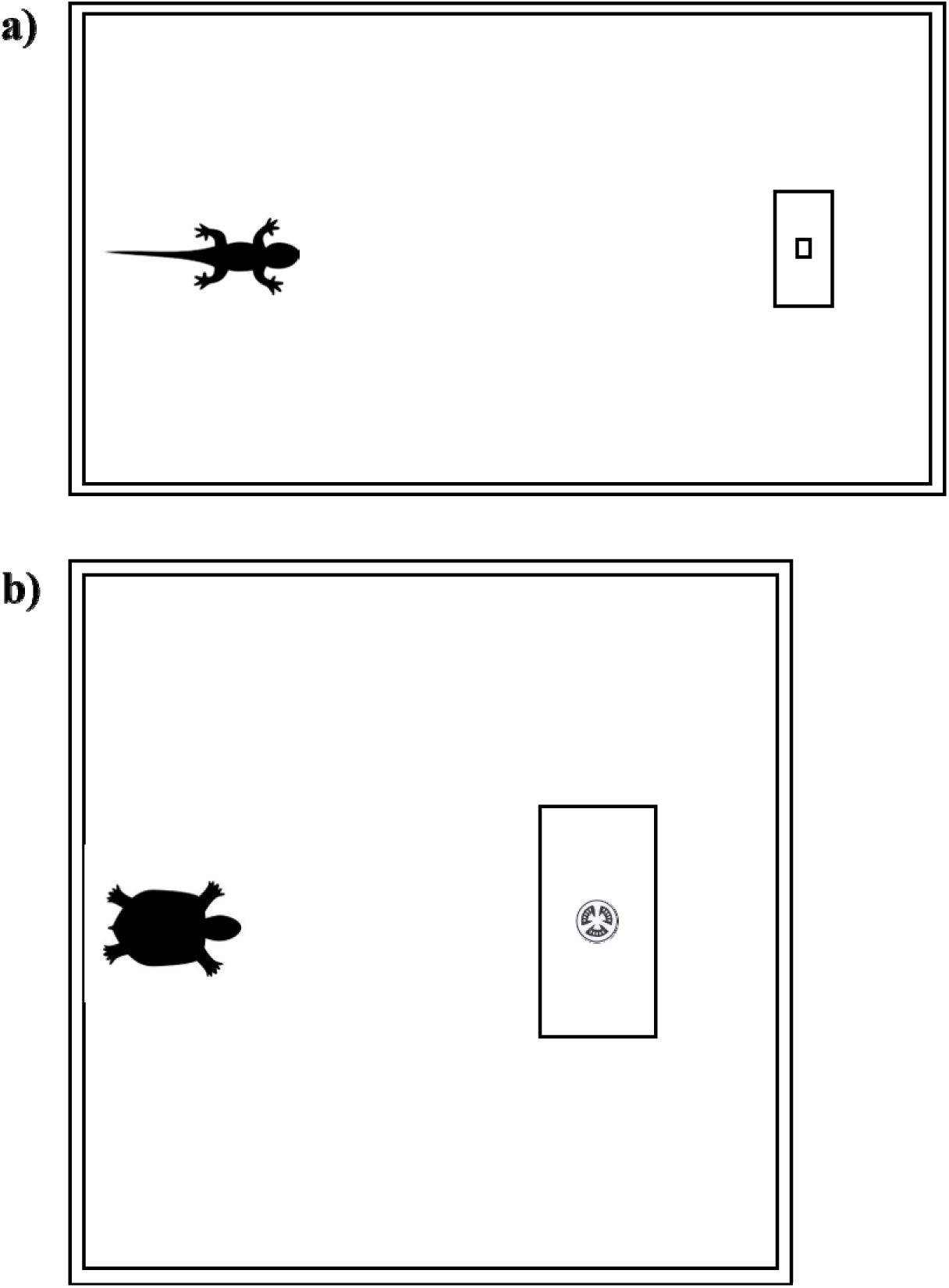
View from above of the apparatuses and transparent cylinders used for bearded dragons (a) and tortoises (b).

### Procedure

#### Training phase

This phase lasted two days during which reptiles learnt to obtain food inside the opaque cylinder. The subjects performed five trials per day with a 30-minute interval between trials. Reptiles had 30 min to reach the food in the opaque cylinder; otherwise, the trial was considered null and was repeated. However, if the subject entered the cylinder, we waited 5 min before removing the cylinder to allow the subject to consume the food.

#### Test phase

The test phase was similar to the training phase, but the opaque cylinder was replaced with the transparent one. In this phase, all reptiles received 20 trials divided into five trials per day for 4 consecutive days with a 30-minute interval between trials. Based on the video recordings, we scored both the reptiles’ accuracy and the time required to enter the cylinder in each trial. For the accuracy, we scored a correct trial when the subject entered the cylinder directly via the lateral openings without touching the cylinder. Instead, we scored an incorrect trial when the subject touched the transparent material before entering the cylinder. To score the time to solve the task, we measured the time from when the subject was inserted in the apparatus to when it entered the cylinder. One-third of the videos of each species were analysed by two different experimenters to assess interrater reliability. Trial-by-trial interobserver agreement was calculated by dividing the number of trials in which experimenters agreed by the total number of trials and then converting the result to a percentage; the mean agreement for total performance was 99%. Even the time to solve the task indicated high interrater reliability (Spearman rank correlation: ρ = 0.944, *p* < 0.001). Therefore, we conducted all the analyses with the database of the first experimenter.

### Data analysis

Analyses were performed in R version 4.3.2 (The R Foundation for Statistical Computing, Vienna, Austria, http://www.r-project.org). Analyses were performed on the frequency of correct trials and were similar for the two species. We analysed the subjects’ accuracy in each trial (correct or incorrect) with generalized linear mixed-effects models for binomial response distributions (GLMMs, ‘glmer’ function of the ‘lme4’ R package). To assess whether accuracy increased day after day and trial after trial (within the same day), we fitted the models with the day (1 – 4) and trial number (1 – 5) as fixed effects. Moreover, we also fitted the time to reach the food (after log transformation) as fixed effect whereas individual ID was fitted as a random effect. Only for tortoises (due to the balanced sample size), we also included sex as a fixed factor.

A Pearson correlation test was used to assess the correlation between the proportion of correct choices and a ‘lateralization’ index calculated as the proportion of entering the cylinder from the preferred side. A lateralization index of 1 means that a subject always entered the cylinder from the same side (always right or always left); instead, a lateralization index of 0.5 means that half of the time the subject entered from the right side and the other half from the left side suggesting no lateralization.

To compare the training phase of the two species, we analysed the subjects’ time to enter the opaque cylinder with a linear mixed-effects model (LMM, ‘lmer’ function of the ‘lme4’ R package). We fitted the model with the species and the day (1 – 4) as fixed effects and individual ID nested within species as a random effect. To compare the accuracy of the two species in the test phase, we fitted a GLMM with the species and the day (1 – 4) as fixed effects and individual ID nested within species as a random effect.

## RESULTS

### Pogona vitticeps

With the opaque cylinder, bearded dragons entered the cylinder in 98 ± 16% of trials (mean ± SD). The mean time required to enter the cylinder was 282 ± 362 seconds (mean ± SD).

With the transparent cylinder, bearded dragons performed 41.25 ± 28.26% correct trials (mean ± SD) in which they retrieved food without touching the transparent cylinder (correct trials). The mean time required to complete a trial was 145 ± 106 seconds (mean ± SD). Performance of the first 10 trials (the measure adopted in other studies [6,10]) was 31 ± 28% correct trials.

The GLMM revealed a significant effect of the time on subjects’ accuracy (χ^2^_1_ = 4.189, *p* < 0.05): the longer the time to reach the food, the lower the performance. Also, both day and trial had a significant effect on subjects’ accuracy (day: χ^2^_1_ = 6.040, *p* < 0.05, Figure 2; trial: χ^2^_1_ = 13.322, *p* < 0.001): bearded dragons’ performances increased day after day and trial after trial (within the day). No interaction was statistically significant (all *p*-values > 0.201).

**Figure 2.**
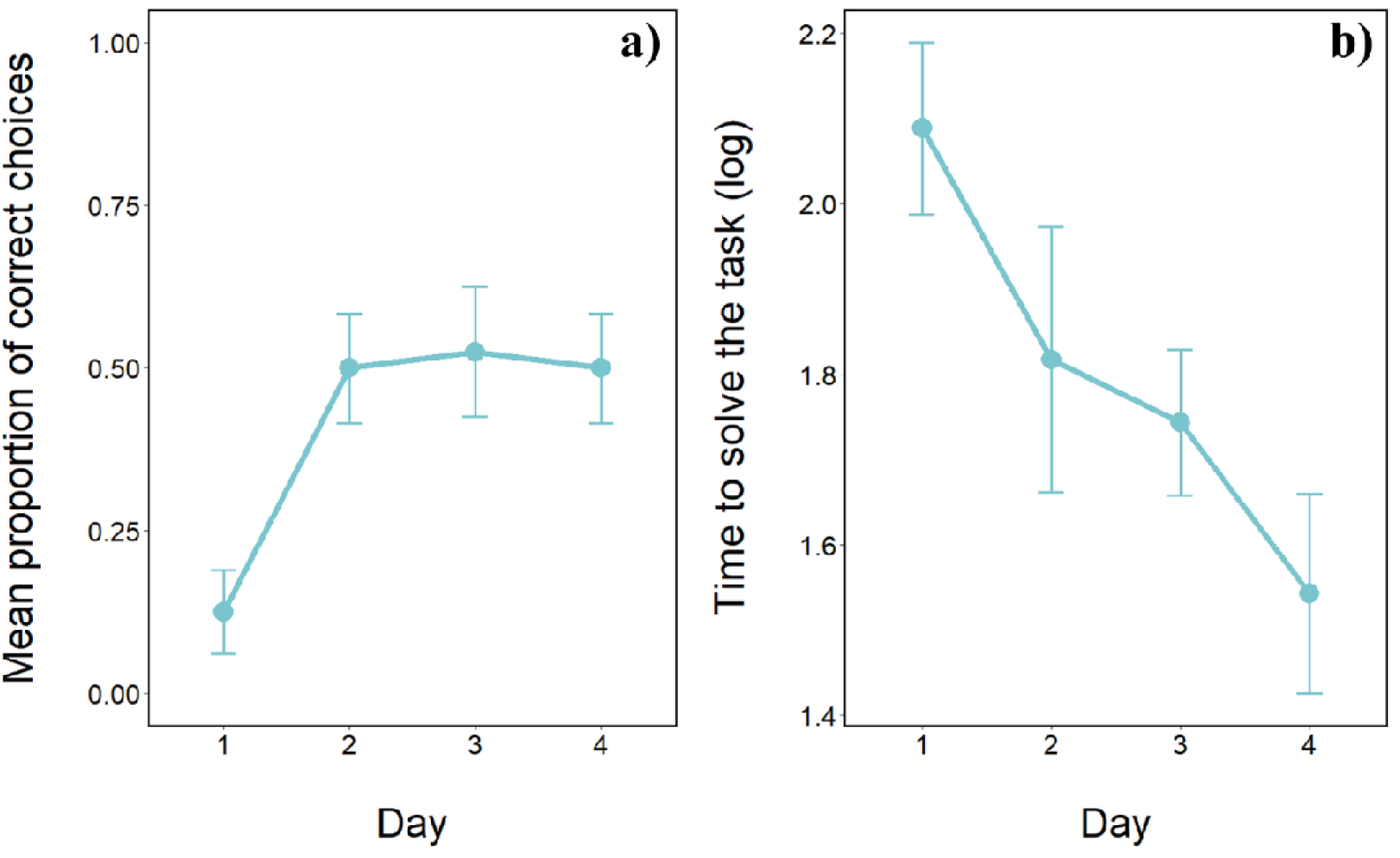
Performance of bearded dragons in the cylinder task. (a) Mean proportion of correct trials in which bearded dragons did not contact the cylinder (mean□±□SE), and (b) time to solve the task (mean□±□SE) over the four days of the test phase.

At the population level there was no left-right bias in the direction bearded dragons entered in the cylinder (mean ± SD: 49.38 ± 20.43 %; one-sample *t* test: *t*_7_□=□-0.087, *p* = 0.934). However, individuals varied considerably in the degree of lateralization and Pearson correlation revealed a positive significant correlation between bearded dragons’ accuracy and lateralization (*r*_6_ = 0.898, *p* < 0.01) suggesting that more lateralized subjects had an advantage in solving the task and reaching the food.

### Testudo hermanni

With the opaque cylinder, male tortoises entered the cylinder in 98 ± 16% of trials (mean ± SD). Their mean time required to enter the cylinder was 201 ± 392 seconds (mean ± SD). Instead, female tortoises entered the cylinder in 98 ± 20% of trials (mean ± SD). Their mean time required to enter the cylinder was 302 ± 494 seconds (mean ± SD). We found no significant difference between the sexes in the time to enter the cylinder (*F*_1,150_ = 0.081, *p* = 0.777).

With the transparent cylinder, tortoises performed 28.13 ± 27.47% correct trials (mean ± SD; females: 34.17 ± 29.16%; males: 22.08 ± 24.49%) in which they retrieved food without touching the transparent cylinder (correct trials). The mean time required to complete a trial was 106.61 ± 137.46 seconds (mean ± SD; females: 145 ± 184; males: 68 ± 35). These data seem to suggest that females have higher performances in terms of correct choices but take longer to complete the task and reach the food inside the cylinder. Performance of the first 10 trials was 22 ± 19% correct trials.

The GLMM revealed a significant effect of the time on subjects’ accuracy (χ^2^_1_ = 26.358, *p* < 0.001): the longer the time to reach the food, the lower the performance. Moreover, females’ performance was higher compared to males’ performance (χ^2^_1_ = 3.957, *p* < 0.05; Figure 3). Also, day had a significant effect on subjects’ accuracy (χ^2^_1_ = 6.737, *p* < 0.01; Figure 3) but not trial (χ^2^_1_ = 2.248, *p* = 0.134): tortoises’ performances increased day after day but not trial after trial (within the day). No interaction was statistically significant (all *p*-values > 0.101) with the exception of time × day (χ^2^_1_ = 9.808, *p* < 0.01): given the same amount of time needed to complete the task, day after day, tortoises’ performance was higher.

**Figure 3.**
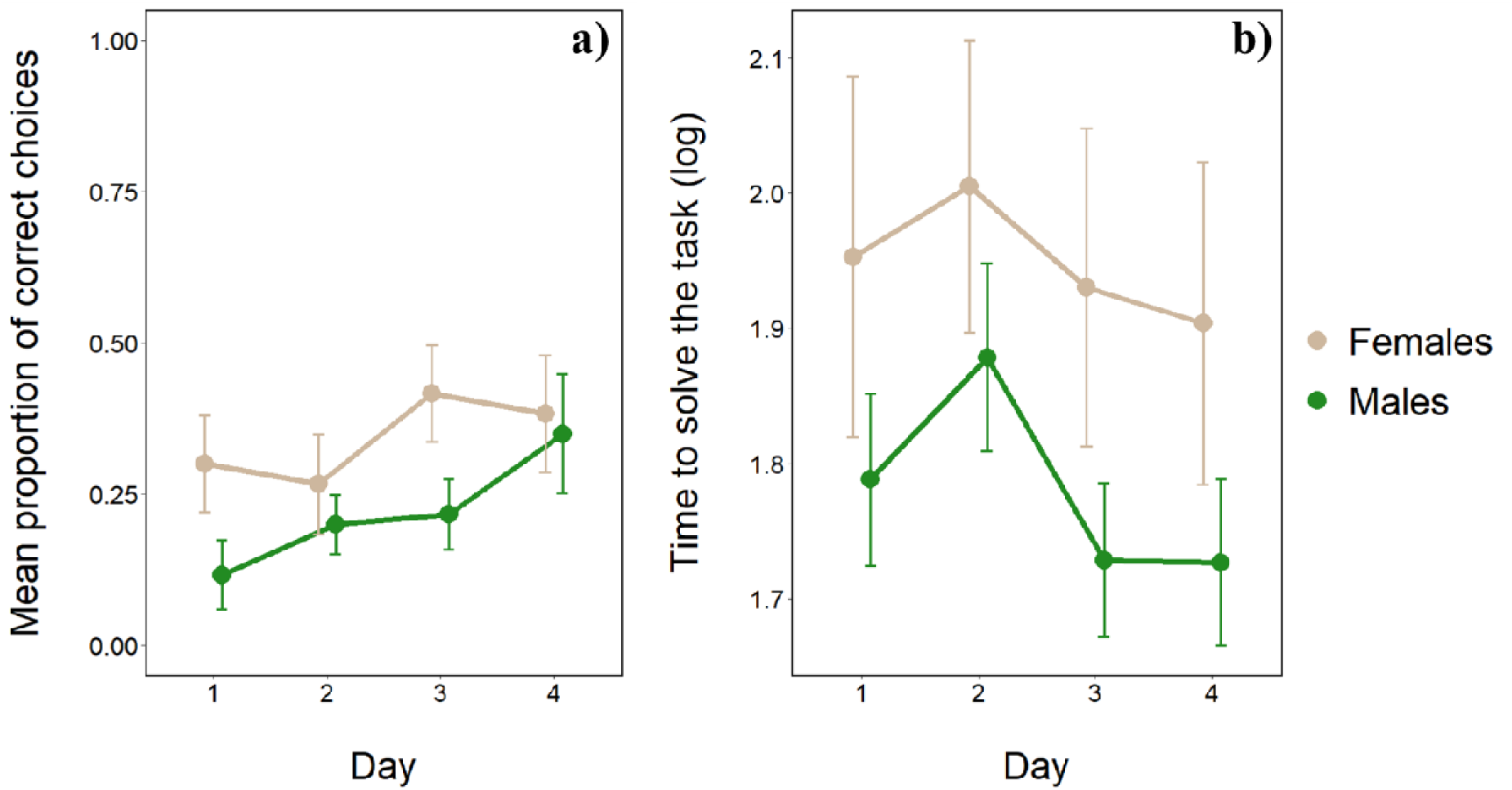
Comparison of the performances of females’ and males’ tortoises in the cylinder task. (a) Mean proportion of correct trials in which tortoises did not contact the cylinder (mean□±□SE), and (b) time to solve the task (mean□±□SE) over the four days of the test phase.

At the population level there was no left-right bias in the direction tortoises entered in the cylinder (males: mean ± SD = 42.50 ± 17.37 %, one-sample *t* test: *t*_11_□=□-1.494, *p* = 0.163; females: 45.00 ± 25.55 %, *t*_11_□=□-0.736, *p* = 0.477; all individuals: 43.75 ± 20.28 %; *t*_23_□=□- 1.510, *p* = 0.145). A large variation in the degree of lateralization was observed also in tortoises and the Pearson correlation revealed a positive significant correlation between accuracy and lateralization (*r*_22_ = 0.405, *p* < 0.05) suggesting that, even for tortoises, more lateralized subjects had an advantage in solving the task and reaching the food.

### Interspecific comparison

When comparing tortoises’ and bearded dragons’ performance with the opaque cylinder, the LMM revealed that bearded dragons took significantly longer to enter in the cylinder (*F*_1,285_ = 6.753, *p* < 0.01). Moreover, performances did significantly change between the two days (*F*_1,283_ = 0.0001, *p* = 0.992) but there is significant difference between the curves of the two species (*F*_1,283_ = 0.037, *p* < 0.01) with tortoises that took significantly less time in the second day whereas bearded dragons did not improve in the time to enter the opaque cylinder.

When comparing tortoises’ and bearded dragons’ accuracy with the transparent cylinder, the GLMM revealed that bearded dragons had a significant higher proportion of correct choices (χ^2^_1_ = 5.319, *p* < 0.05; Figure 4). Moreover, performances significantly increased day after day (χ^2^_1_ = 19.843, *p* < 0.001; Figure 4) with no significant difference between the learning curves of the two species (χ^2^_1_ = 0.808, *p* = 0.369; Figure 4).

**Figure 4.**
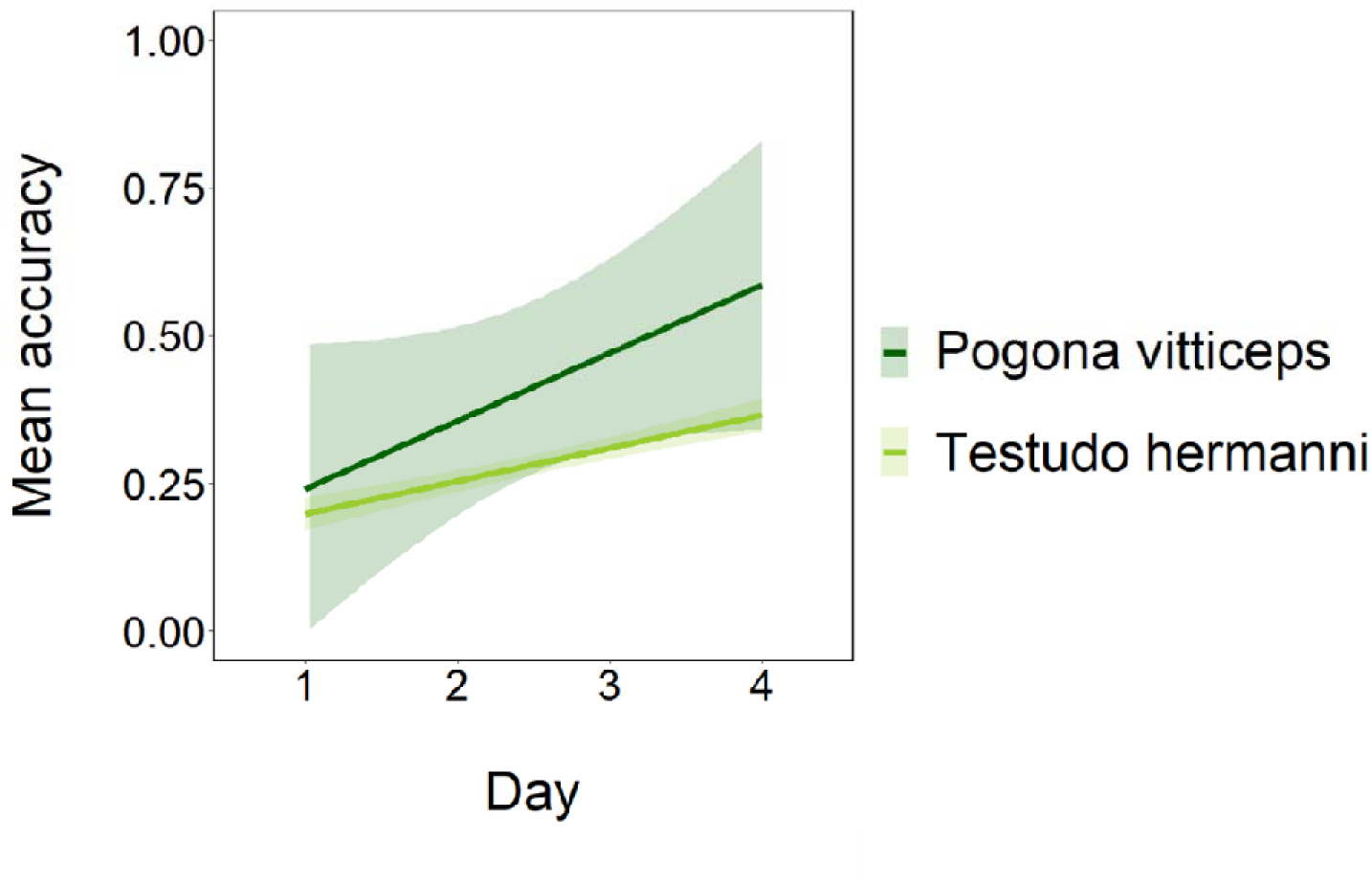
Comparison of mean accuracy between the two species. The shaded areas represent 95% confidence intervals around the estimated effects.

## DISCUSSION

Our study provides the first data on inhibitory capacity in reptiles that can be used for direct comparisons with other vertebrates studied so far. The procedure used in our study, the transparent cylinder test, represents the gold standard for measuring inhibitory control in animal species [6,7,34]. In a preliminary phase, an animal is trained to retrieve food from an opaque cylinder, which is replaced with a transparent one during the trials of the testing phase. This introduces a conflict between the correct motor response learned during training trials and the direct sight of the food reward, which attracts the animal and compels it to reach for the food passing through the barrier.

Both *Testudo hermanni* and *Pogona vitticeps* showed some capacity to inhibit a prepotent but ineffective response, and to temporarily move away from the goal to obtain the reward. As observed in the other vertebrates, performances increased day after day but in both reptiles, even after 4 days and 20 trials, most individuals performed more than 50% of the trials incorrectly. Tortoises and bearded dragons showed a similar improvement rate in the course of the experiment but overall, bearded dragons made a significantly higher percentage of correct trials. Notably, when considering the time required to enter the opaque cylinder during the training phase, tortoises, on average, needed less time than bearded dragons to reach the food and showed a more pronounced improvement over the course of training. This suggests that the differences in performance likely were not due to variations in learning abilities.

How do the inhibitory capacities of reptiles compare to those of other vertebrates? Studies conducted with the same paradigm disclosed the existence of a large interspecific variation in inhibitory capacities both between and within vertebrate classes [6,7]. Species with large complex brains, such as apes and corvids, show very high percentages of correct responses, in some cases approaching 100% (Figure 5). In contrast, small passerine birds and micromammals show much lower percentages, ranging between 25% and 40% (Figure 5). As for teleost fish, the guppy (*Poecilia reticulata*) which has been extensively studied for inhibition capacities [10,35–37], shows a performance of 50% in the transparent cylinder test, far below that of large-brained species but higher than almost all bird species and some mammals. There is some evidence that inhibition capacities may be similar or even greater in other teleosts [38,39].

**Figure 5.**
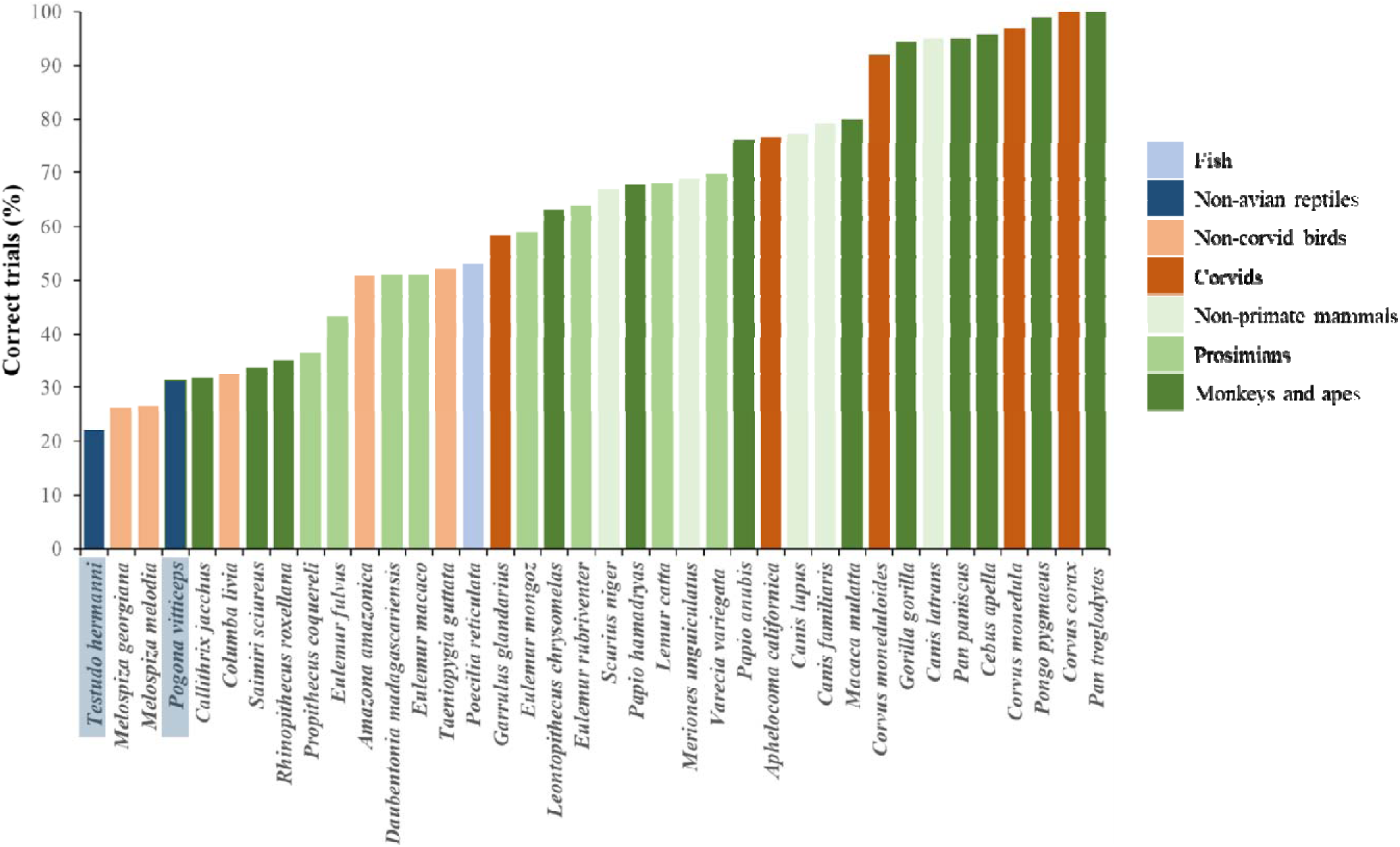
Comparison between the performance of Hermann’s tortoises and bearded dragons in the cylinder task (black bars) and that of different mammalian and avian species tested in the same task by MacLean et al. [6] and Kabadayi et al. [7], and fish tested by Lucon-Xiccato et al. [10]. Bars represent the mean percentage of correct trials. We used the performance of reptiles in the initial 10 trials to allow the comparison with the other species.

When we analyse the performance of the first 10 trials (the measure adopted in the other studies [6,11], both reptiles performed poorly compared with other species. *P. vitticeps* achieved 31% correct responses, placing it at the lower end of the range observed across approximately 50 vertebrate species and well below the performance of teleost fish. *T. hermanni* performed even worse, achieving only 22% correct trials.

Inhibitory capacities have also been examined in scincid lizards using a task similar to the transparent cylinder test [18,40]. The five skink species tested showed an average performance of 34.6% correct responses, with interspecific variation ranging from 26% to 45%. These values were below those observed in teleosts and most homeotherms. However, the experiments employed a different method, measuring detour ability around a perforated wire mesh barrier. The barrier’s characteristics strongly influence performance in detour tasks [19–21], and tests using wire mesh barriers tend to significantly overestimate inhibitory capacity. Therefore, results obtained on skinks cannot be directly compared with data from other species

Currently, we can only speculate about the possible explanations for the difference between the two species of this study and about the reasons reptiles exhibit poorer inhibitory control abilities compared to warm-blooded vertebrates. The comprehensive comparative study conducted on over thirty species of endotherms suggests that absolute brain size is the primary predictor of these differences in inhibitory capacity [6]. In reptiles, both absolute and relative brain sizes are much smaller than in homeotherms [8,41]. In this context, the lower performance of our two species compared to most mammalian and avian species aligns with the hypothesis of a positive relationship between inhibitory capacity and absolute brain size. Moreover, as neuron density increases with brain size across vertebrates, species with a higher encephalization index, such as mammals and birds, possess disproportionately larger numbers of neurons compared to species with smaller relative brain sizes, such as reptiles [8]. Therefore, the performance of the reptilian species examined so far could also support the hypothesis of a relationship between inhibitory capacity and neuron count. Other factors could be at play. For instance, the method of accessing food or other targets may influence inhibition; species using hands or paws, like primates, may exert better inhibitory control than those relying on direct access with the mouth, as manual manipulation allows for a more controlled approach [6]. Clearly, more data from additional reptilian species, especially larger reptiles, are needed before determining whether the hypotheses proposed for homeotherms can be extended to all amniotes.

The exception of teleost fish remains unresolved. These species have brains some orders of magnitude smaller than warm-blooded vertebrates, and significantly smaller than the reptiles studied [41], yet they overperform various mammals, most birds, and all the reptiles examined so far in the cylinder test. A possible explanation for the exception represented by fish may lie in the unique evolutionary history of the bony fish genome and cognition. Over the last decade, it has been discovered that teleost fish possess several cognitive abilities once thought to be exclusive to land vertebrates. For example, they are capable of cooperating to achieve shared goals, acquiring new foraging and antipredator behaviour s through social learning, using tools, engaging in sophisticated parental care, and even demonstrating behaviours that could be classified as cultural traditions (reviewed in [16,42,43]). It is now hypothesized that the evolution of such advanced cognitive abilities is linked to a whole-genome duplication event that occurred over 400 million years ago, after the teleost fish diverged from the lineage leading to terrestrial vertebrates. This duplication freed many genes to evolve new functions, which may explain both the remarkable biological diversity seen in modern fish and the development of complex cognitive abilities in small-brained organisms [17,44]. Other hypotheses have been proposed to explain this paradox. One possibility is that the general organization of their nervous system qualitatively differs from that of tetrapods, for example in glia/neuron ratios or neuronal density [12]. It was also suggested that fish may rival endotherms in certain cognitive abilities but lack entirely other functions - particularly domain-general processes – that are computationally demanding [45].

Notably, the disparity observed between tortoises and bearded dragons is challenging to explain through brain size or neuron count hypotheses. If differences in brain size or neuron number exist, they would likely favour tortoises, suggesting that in this case other factors are involved [8,37]. For instance, an effective inhibitory control system may have evolved in some species but not in others due to ecological factors and the specific ways in which animals interact with their environment. For example, MacLean and collaborators [6] found that, in addition to brain size, inhibitory control abilities positively correlate with diet breadth. They suggested that a more diverse and complex diet could drive the evolution of larger brains and advanced cognitive functions, which in turn may have facilitated the emergence of more sophisticated inhibitory capacities [6]. Alternatively, diet type may have directly influenced the evolution of more efficient inhibitory abilities, as species with broader diets face complex decision-making in diverse foraging contexts. This ecological factor could explain the interspecific difference observed in our study, given that *P. vitticeps* is a dietary generalist, while *T. hermanni* is almost exclusively herbivorous [46–48]. Finally, both individual and species differences may be mediated by non-cognitive factors, such as feeding motivation and body condition [49], or prior experience with transparent barriers [50].

Motivation has been found to influence performance on cognitive tasks [51–52]. In tests measuring inhibitory control, some species exhibit a decline in performance as motivation increases [35,49]. However, making comparisons between phylogenetically distant species is extremely challenging, as they can differ significantly in physiology, energy requirements, diet, and ecology. Using the time taken to reach the food during the training phase as a proxy for feeding motivation, we found that tortoises were faster than bearded dragons, which contradicts the expectations based on the feeding motivation hypothesis. As for prior experience with transparent barriers, its effect has been observed in some species but not in others [36,50,54]. In our study, although both species had some level of exposure to transparent or semi-transparent barriers (e.g., wire mesh), the bearded dragons likely benefited from more frequent prior exposure to such barriers. In contrast, tortoises, which are known to encounter difficulties with transparent surfaces, tend to rely more on physical strength to navigate obstacles in their natural environment. For instance, research indicates that tortoises in dense vegetation environments primarily use force to push through obstacles [55]. Moreover, when confronted with physical barriers, such as steep steps, tortoises do not attempt sophisticated detour routes but rather persistently use their body strength to overcome these barriers [56]. This reliance on force over detour may also apply to artificial transparent barriers: tortoises may not interpret transparent obstacles as true barriers, resulting in behaviour focused on direct physical engagement rather than detouring or inhibiting prepotent responses.

Cognitive sex differences have been reported in many vertebrates. These differences span a wide variety of cognitive functions, including numerical abilities, spatial cognition, problem-solving skills, etc. [57–62]. With the exception of spatial abilities, where males generally outperform females, findings across other cognitive domains indicate that either sex may excel depending on the function and species studied.

Recent studies have also reported sex differences in executive functions, a particular class of advanced cognitive processes, including cognitive flexibility, working memory, and inhibitory control, that enable the control of goal-directed behaviour [63–66]. Specifically, regarding inhibitory control, sex differences have been documented across a range of mammals, fish, and birds, sometimes favouring females and other times favouring males (reviewed in [59]). In this study, we found that sexual differences in executive functions exist also in reptiles. In the only species, *T. hermanni*, for which the sex ratio and sample size allowed for such an analysis, we found that females outperformed males in the cylinder test. The temporal pattern of performance suggests that this difference is not due to learning differences, as females showed superior performance from the beginning of the test, and both sexes improved at a similar rate over time.

Cognitive sex differences are generally thought to arise from differential selection pressures driven by ecological sex differences or by varying mating and reproductive roles. However, it is often challenging to pinpoint the exact cause of such differences in a species unless its life history, mating system, and any sex differences in foraging or other ecological aspects are known in detail. As far as we know from *T. hermanni*’s biology, there are no significant trophic niche differences between the males and females, and the species’ mating system does not differ markedly from other vertebrates with a promiscuous mating system and no parental care [47,67,68].

Sex differences in cognitive tests may also stem from non-cognitive factors. Although the procedure was identical for both sexes, motivation could differ. For instance, field studies show that females invest more in reproduction due to the energy costs of egg production, resulting in poorer body condition during the breeding season, reduced annual survival, and shorter lifespan compared to males [69,70]. However, in captivity, where food is provided ad libitum and foraging can occur continuously, it is unclear whether this would lead to different feeding motivations. In any case, the direction of this effect would contradict the relationship between motivation and performance observed in other species [35].

Another important factor could be body size, which affects brain volume. In wild populations, *T. hermanni*, males are smaller than females. This is partly compensated by the relatively larger size of male heads although it remains unclear whether a relatively larger head also corresponds to a larger brain [65]. In our study, we tried to match body size between the sexes as much as possible, and indeed, there were no significant differences in size between males and females.

An interesting finding of this study is the positive correlation, observed in both species, between an individual’s degree of lateralization and its performance in the cylinder test. The nature of this relationship remains unclear. Recent debates have centered on the evolutionary advantages of brain lateralization, which may confer adaptive benefits by enabling the simultaneous processing of different types of information in the two hemispheres [23,25,27]. This ability could help animals manage limited attention by performing two cognitive tasks at once, for instance feeding or mating while remaining vigilant for predators. In our case, it is difficult to argue that the experimental situation required sharing attention between two simultaneous tasks unless we assume that being in an unfamiliar environment outside of their home cage triggered automatic vigilance for potential predators. There are, nonetheless, several other cases like this where it seems challenging to invoke the advantage of more efficient multitasking to explain the superior performance of lateralized animals [23,71] suggesting that other advantages must contribute to explain the ubiquity of lateralization. Another classic hypothesis for the ubiquity of lateralization is that it helps avoid the simultaneous activation of incompatible responses that might arise from symmetrical duplication of brain areas in both hemispheres. One situation where this might be relevant is when an animal needs to move around an obstacle to reach a goal. Since detouring to the left or right are functionally equivalent responses, a perfectly symmetrical nervous system could lead to simultaneous activation or mutual inhibition, potentially interfering with the execution of the response or at least slowing the decision-making process. In this sense, the influence of lateralization may not be on inhibitory control per se, but rather on the execution of the specific task in the transparent cylinder test. To test this hypothesis, one would need to assess inhibitory control using a task that does not involve spatial right-left components.

In conclusion, as with warm-blooded vertebrates, reptiles appear capable of inhibiting prepotent but inappropriate responses to a given context. By employing the same test used across a wide range of vertebrates, we were able to compare the inhibitory control of reptiles directly with that of other species. Our findings show that the inhibitory control abilities of reptiles, as measured by the transparent cylinder test, are lower than those of other amniotes observed so far. These results align with the prevailing hypothesis that the size and complexity of the nervous system are the primary predictors of inhibitory control abilities. However, the existence of significant exceptions to this rule among vertebrates suggests that species-specific ecological factors or non-cognitive variables may also play an important role in determining interspecific differences. This study confirms previous findings in other species, demonstrating that sex differences in inhibitory capacities also exist in non-avian reptiles and highlights for the first time that hemispheric lateralization can enhance cognitive functions, including inhibitory control, in cold-blooded amniotes.

## ADDITIONAL INFORMATION

## Acknowledgments

We wish to thank Tommaso Pecchia for his help in building and setting up the experimental apparatus and materials used with tortoises.

## Data availability statement

The authors confirm that the data supporting the findings of this study are available within the article’s supplementary materials.

## AUTHOR CONTRIBUTIONS

This study was conceptualized by A.B., who laid the groundwork for the research. The design and implementation of the study on tortoises were carried out with the guidance of V.A.S. and G.S., while the study on bearded dragons was developed in collaboration with A.W. The experiments were performed by M.S., who codified and analysed row data. The original draft of the manuscript was prepared by M.S. and A.B. All the authors read, revised, and approved the final draft of the manuscript.

